# How much does it take to be old? Modelling the time since the last harvesting to infer the distribution of overmature forests in France

**DOI:** 10.1101/2021.02.08.430252

**Authors:** Lucie Thompson, Eugénie Cateau, Nicolas Debaive, Frédéric Bray, André Torre, Patrick Vallet, Yoan Paillet

## Abstract

**Aim:** The distribution of overmature forests in metropolitan France is poorly known, with only a few well-studied prominent sites, and has never been evaluated countrywide. Here, we modelled French forest reserves’ time since the last harvesting operation - a proxy for forest maturity - then inferred the current statistical distribution of overmature forests (i.e. forests over 50 years without harvesting) in France.

**Location:** Metropolitan France

**Methods:** We used inventories from forest reserves and managed forests to calibrate a generalised linear mixed model explaining the time since the last harvesting with selected structural attributes and environmental variables. We then projected this model on the independent National Forest Inventory dataset. We thus obtained an updated estimation of the proportion and a rough distribution of overmature forest stands in metropolitan France.

**Results:** We found that high basal area of very large trees, high volumes of standing and downed deadwood, high diversity of tree-related microhabitats and more marginally diversity of decay stages best characterized the time since the last harvesting. Volumes of stumps and high density of coppices translating legacy of past forest management also distinguished more overmature plots. Our projection yielded an estimated 3% of French forests over 50 years without harvesting mostly located in more inaccessible areas (i.e. mountainous areas) and a promising proportion of future overmature forests if left unharvested.

**Main conclusions:** Our study showed that the time since the last harvesting is a good proxy for a combination of stand structure attributes key in characterising overmature temperate forests. It gives the first robust statistical estimate of the proportion of overmature forests and may serve to report on their status in metropolitan France. Our method could be implemented at a larger spatial scale, notably in countries with accessible National Forest Inventory and calibration data, to produce indicators at international level.

## 1 INTRODUCTION

Old-growth forests have a key role in the mitigation of climate change. They act as carbon storage and sinks (Achard & Hansen, 2012; Frey et al., 2016; Luyssaert et al., 2008). They have a role in the protection of water resources and the prevention of soil erosion (Brockerhoff et al., 2017; Watson et al., 2018) and also host a myriad of features with high conservation value promoting biodiversity (Bauhus et al., 2009; Burrascano et al., 2013; Larrieu et al., 2018; Paillet et al., 2010, 2015, 2017).

Wirth et al., (2009) present various ways to define forest maturity. Their structural definition is a combination of dominant tree species’ age and estimated longevity, with stands considered old-growth if they harbour dominant species older than half of their longevity. In practice, this definition is problematic in that (1) tree age is best determined by core sampling, a tedious task which may underestimate tree age (Speer, 2009) and (2) longevity is not a well-known species feature, it depends on various factors and might be biased by the long history of forest management displayed in European forests, resulting in most trees being harvested before they die of natural senescence (Cateau et al., 2015). To work around those issues, international reporting generally uses the time since the last harvesting as a proxy for forest maturity or naturalness (see indicator 4.3 in the State of Europe’s forests, (Forest Europe, 2020). By “harvesting” we considered any human intervention extracting wood biomass from the forest, including various harvesting intensity and management regime, ranging anywhere from thinning to final cut, and from strict forest reserves to regularly harvested forests. Indeed, whatever their degree of maturity, forests bear the legacy of past – sometimes intensive – forest use in their stand structure, notably silvicultural treatments such as coppice-with-standards (Lassauce et al., 2012). The use of the time since the last harvesting is a way of assessing the degree of naturalness of the forest in question, with increased occurrence of maturity feature or structural attributes in older unmanaged plots. Those plots display more trees with larger diameters at breast height (Burrascano et al., 2013; Heiri et al., 2009; Paillet et al., 2015), a higher abundance of tree-related microhabitats (Larrieu et al., 2018; Paillet et al., 2017; Winter & Möller, 2008) along with high volumes of deadwood (Harmon, 2009; Siitonen et al., 2000). Those specific structural attributes harboured by more mature forests also have high conservation value for biodiversity (Bauhus et al., 2009; Burrascano et al., 2013; Siitonen et al., 2000).

In France, it is acknowledged that 50 years of abandonment are a minimum for a forest to display sufficient maturity features (MAAF & IGN, 2016). Such a threshold may not qualify a given forest as “old-growth”, but still represents a transition phase towards this state. We herein call these forests “overmature” (Wirth et al., 2009) and distinguish them from primary forests which, according to the Food and Agriculture Organisation are forests with no clearly visible indications of human activities and disturbance (FAO, 2015). However primary forests patches – like any forest with natural succession - can be at various stand succession stages and levels of maturity including overmature and old-growth.

Despite the many services they provide (Paillet et al., 2010; Watson et al., 2018) overmature and primary forests remain scarce at the global scale, especially in western Europe and most are, as yet, unprotected (Forest Europe, 2015; Sabatini et al., 2018; Sabatini, Keeton, et al., 2020). In their report for the European Commission, Barredo et al., (2021) assessed that primary and overmature forests constituted 2.4% of forested area in Europe. There is a recognised need for a global map of old-growth and primary forest patches (Barredo et al., 2021; Chiarucci & Piovesan, 2020) and the “EU Biodiversity Strategy for 2030” explicitly mentions that “it will be crucial to define, map, monitor and strictly protect all the EU’s remaining primary and old-growth forests” notably by designating at least 10% of Europe’s land to strict protection (European Commission, 2020). Indeed, the restoration of degraded forest is often more costly than to conserve existing ecosystems, and forest degradation can only be partially reversed on a reasonable time scale (Chazdon, 2008). However, studies of high conservation value forests such as overmature forests in France have targeted, to date, only a few emblematic reserves (Christensen et al., 2005; Mountford, 2002; Pontailler et al., 1997). (Sabatini, Keeton, et al., 2020) evaluated the area of primary forests in France to less than 0.1% of the forested area and the sole estimation for the proportion of overmature forest stands in France dates back to 1993 and was evaluated to about 3% (MAAPRAT-IFN, 2011). This estimation has not been updated since and relied more on expertise rather than sound data or rigorous analytical approach. There is therefore a knowledge gap concerning the proportion, distribution and overall characterisation of overmature forests that needs to be filled.

In this study, we used inventories of forest stand structure issued from forest reserves and managed forests to calibrate a generalised linear mixed model explaining the time since the last harvesting with selected attributes of maturity, herein called structural variables, combined with environmental variables. We hypothesized that time since the last harvesting would be positively influenced by a high volume of large logs and snags (Heiri et al., 2009; Portier et al., 2020; Siitonen et al., 2000) and high basal area of very large trees (Burrascano et al., 2013; Paillet et al., 2015) a high diversity of tree-related microhabitats (Larrieu et al., 2018; Paillet et al., 2017; Winter & Möller, 2008) and decay stages (Siitonen et al., 2000; Winter & Möller, 2008; Wirth et al., 2009). We expected time since last harvesting to be negatively correlated with the volume of stumps (Paillet et al., 2015; Siitonen et al., 2000) and stem density of medium trees (Paillet et al., 2015). We also tested other structural variables that would bear witness to past management (e.g. proportion of coppice). Finally, we expected overmature forests to be in more remote and less productive areas (Levers et al., 2014, 2018; Sabatini et al., 2018), hence positively correlated with elevation or slope and soil fertility.

We projected this model on an independent nation-wide dataset issued from the National Forest Inventory. Thus, we obtained an updated estimation of the proportion and a rough distribution of overmature forest stands (i.e. abandoned for over 50 years) in metropolitan France. This study may serve to report on the status of overmature forests in France, as well as a guideline for inferring their distribution in other North-western European countries using maturity features in similar temperate forest types.

## 2 METHODS

### 2.1 Training data

We worked with a dataset issued from a monitoring program that has been implementing forest stand structure description in French forest reserves since 2005, to monitor the evolution of forest attributes. A stratified sample design encompasses the variability within a forest reserve. Plots can be comprised in strict forests reserves, where harvesting is prohibited, special forest reserves, where management targets specific habitat or species (e.g. forest ponds), as well as managed stands in forests reserves where harvesting may be allowed under certain conditions (Table 1). They also include plots in both private and public forests, but with a vast majority of public forests. One forest reserve can host up to several hundred plots depending on the size of the represented area and its layout. We kept the plots where the time since the last harvesting was recorded by forest managers, by looking into current or old management plans. Plots from this network show a larger abandonment gradient than French forests in general (see Appendix S1 Figure S1.1 in Supporting Information).

**Table 1:** Description of the training dataset (4 728 plots and 71 forest reserves) split by the management status of plots. Special forest reserves include future forest reserves (not yet designated), national and regional forest reserves and Natura 2000 sites

| Plot status | Number of plots | Mean basal area of living trees ($\text{m}^2.\text{ha}^{-1}$) | Mean volume of standing dead trees ($\text{m}^3.\text{ha}^{-1}$) | Mean volume of downed deadwood ($\text{m}^3.\text{ha}^{-1}$) | Mean time since the last harvesting (years) |
| --- | --- | --- | --- | --- | --- |
| **Managed stands in forest reserves** | 228 | 34.9 | 10.6 | 17.4 | 30.8 |
| **Special forest reserves** | 2628 | 27.6 | 14.5 | 28.8 | 31.5 |
| **Strict forest reserves** | 1872 | 28.4 | 10.7 | 20.4 | 33.6 |

4 728 plots in 71 reserves were used for modelling (Figure 1). We only kept the plots with spatial coordinates and classified into one of the six most abundant tree species groups (see below and Appendix S4 for definitions): pure spruce and fir plots accounted for 11% of the total plots, pure beech plots accounted for 14%, mixed broadleaved [over 75% of multispecies broadleaved] 32%, mostly broadleaved [50 – 75% of broadleaved] 8%, mostly conifers [50 – 75% of coniferous] 9% and pure deciduous oaks 7% (Fig.S1.2). As a comparison, overall French production forests is comprised of 65% of broadleaved forests, 21% of coniferous forests and 11% of forests with either mostly broadleaves or mostly coniferous trees [50 - 75 % of basal area] (MAAF & IGN, 2016). Our modelling dataset has therefore slightly less coniferous and broadleaved and more “mostly coniferous” or “mostly broadleaved” forest types than the average French forest.

**Figure 1:**
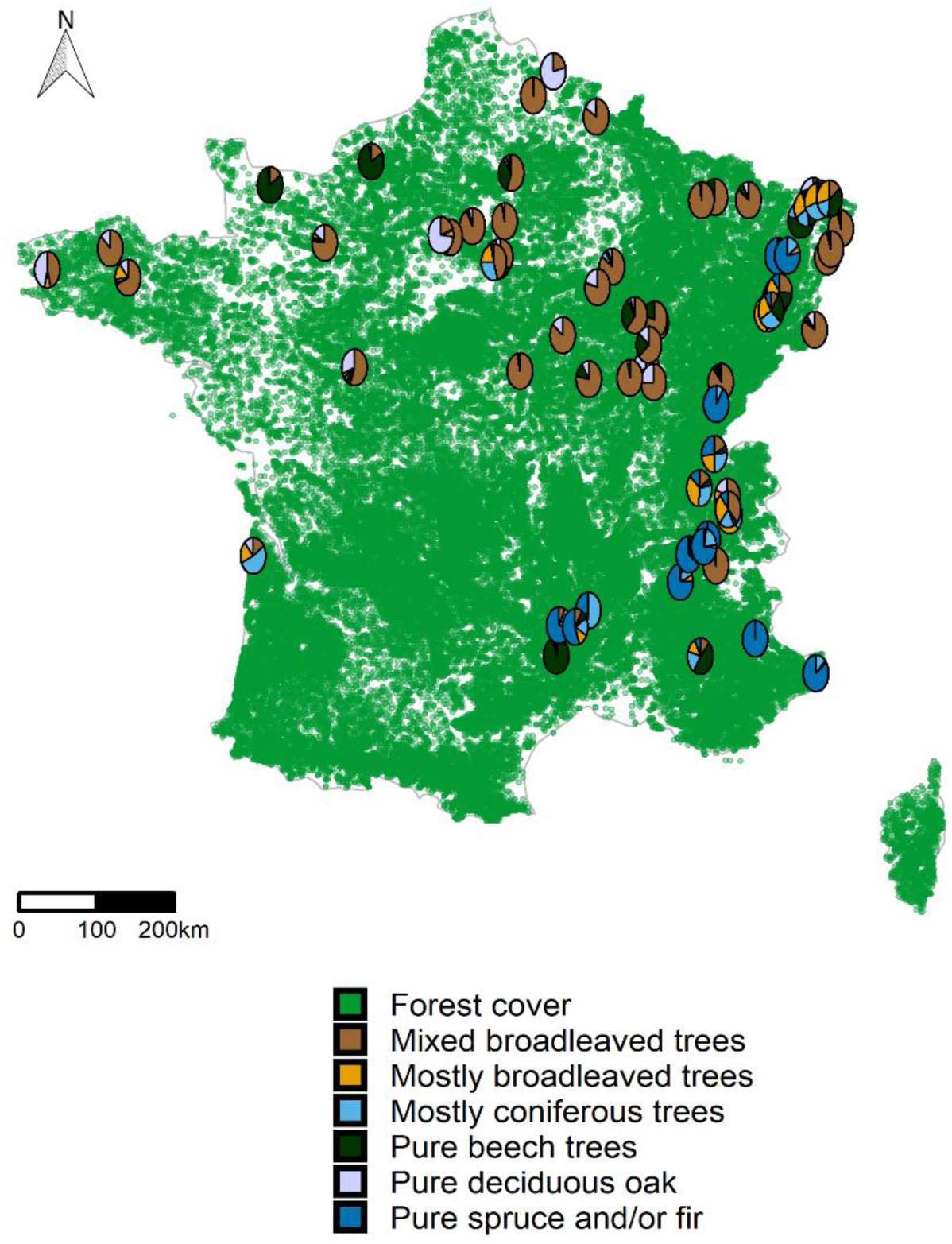
Distribution of 71 reserves from the training data kept for modelling. Each pie plot shows the forest types of the reserve’s constituting plots. The green background shows French forest cover (from the National Forest Inventory).

We did not keep plots over 110 years since the last harvesting as we judged the data too unreliable. We left out reserves with less than 20 remaining plots to ensure sufficient data for the averaging of the random effect (see below). We graphically checked that forests subjected to different management regimes were equivalent in terms of site fertility, precipitation and mean annual temperature, therefore not introducing bias in the dataset (Appendix S1, Figure S1.3).

The protocol for the stand structure surveys is detailed in Appendix S2. Living trees with a diameter at breast height (DBH) under 30 cm were all measured within a fixed 10 m radius plot. Larger trees were measured if they were comprised in a 3% angle count sampling (Paillet et al., 2015). Coppice and standard stems were differentiated. Each shoot from the same stump was measured individually. Standing deadwood under 30 cm and standing and downed deadwood over 30 cm in diameter were measured respectively on a 10 m and 20 m radius plot, discriminating between stumps (under 130 cm tall) snags and standing dead trees. Downed deadwood with a diameter under 30 cm was surveyed on three linear 20 m long transects. Decay stage as well as diameter were recorded off all sampled deadwood (see Appendix S2). Finally, tree-related microhabitat surveys followed the protocol detailed in Paillet et al. (2019) by visually inspecting trees and recording presence of microhabitats. Different microhabitat classifications have been used over the years. Therefore, we created a harmonized classification based on Larrieu et al. (2018)’s microhabitat forms to narrow it down to one homogeneous classification (Appendix S3).

### 2.2 Structural attributes and environmental variables

Among the attributes surveyed, we kept the basal area, volume and stem density of living trees, volume of standing deadwood (stumps, snags and standing dead trees) and volume of downed deadwood (logs) (see Paillet et al., 2015). Diversity of decay stages and tree-related microhabitats were calculated at the plot scale with the Shannon index, one type corresponding to either decay stage associated with tree species and type of deadwood (stump, snag, standing dead tree, downed deadwood) or tree-related microhabitat form associated with species and type of wood (dead or alive) (see Table 2). We split observations into diameter classes: small trees [DBH: 17.5, 27.5cm], medium trees [27.5, 47.5], large trees [47.5, 67.5] and very large trees over 67.5 cm.

**Table 2:** Structural features chosen as candidate variables for modelling the time since the last harvesting of forest plots, and the literature supporting them. “Hypothesis” relates to the expected effect of the structural feature on the time since the last harvesting: « + » (« - ») means we expect a positive (negative) effect of the feature on the time since the last harvesting « ++ » means it is a very common feature found in most articles dealing with forest maturity. This table is based on a literature review, but other attributes were tested considering the local specificities of our dataset.

| Structural features | Variable | Hypothesis | Source |
| --- | --- | --- | --- |
| Living trees | Density and basal area of large and very large trees | ++ | Burrascano et al., 2013; Paillet et al., 2015 |
|  | Density of living trees | + | Portier et al., 2020 |
|  | Total basal area of living trees | + | Heiri et al., 2009 |
|  | Density of medium sized living trees | - | Paillet et al., 2015 |
| Decay stages | Diversity of decay stages | + | Siitonen et al., 2000; Winter & Möller, 2008; Wirth et al., 2009 |
| Tree-related microhabitats | Diversity of tree-related microhabitats | ++ | Larrieu et al., 2018; Paillet et al., 2017; Winter & Möller, 2008 |
| Downed deadwood | Volume of downed deadwood | ++ | Bauhus et al., 2009; Heiri et al., 2009; Paillet et al., 2015; Portier et al., 2020; Siitonen et al., 2000; Vandekerckhove et al., 2009 |
| Standing deadwood | Volume of stumps | - | Paillet et al., 2015; Siitonen et al., 2000 |
|  | Volume of standing dead trees and snags | ++ | Heiri et al., 2009; Portier et al., 2020; Siitonen et al., 2000 |

We extracted environmental variables using plot locations (see Table 3 for the sources):

**Table 3:** Environmental covariates tested for modelling and where they were sourced

| Category | Covariate | Units | Source |
| --- | --- | --- | --- |
| Topographic | Elevation | Meters | BD ALTI v.2.0 : digital terrain model 25 m (IGN) |
|  | Slope | Degrees | Calculated<sup>1</sup> with BD ALTI v.2.0 |
|  | cosine(Aspect) | - | Calculated with BD ALTI v.2.0 |
| Climatic | Mean annual temperature (BIO2) | degrees Celsius | WorldClim v.2 (Fick & Hijmans, 2017) |
|  | Total annual precipitation (BIO12) | Millimetres | WorldClim v.2 |
| Edaphic | Soil pH<sup>2</sup> | - | DIGITALIS dataset - (Laboratoire SILVA, n.d.) (Université de Lorraine-AgroParisTech-INRA) |
|  | C/N ratio<sup>2</sup> | - | DIGITALIS dataset |
|  | Maximum water holding capacity<sup>3</sup> | Millimetres/cm of soil | DIGITALIS dataset |
| Forest type |  | - | Adaptation of the *BD forêt* v.2 (IGN, 2016a) |
| Forested region | SylvoEcoRegion | - | National Forest Inventory (L'IF n°14, 2011) |
| Site index |  | - | (Toïgo et al., 2015) |

<sup>1</sup> « terrain » function from R package *raster* (Hijmans, 2020)
<sup>2</sup> The C/N ratio and soil pH are bioindicated for the first soil layer using floristic surveys from the National Forest Inventory (NFI) and the EcoPlant database (Gégout et al., 2005)
<sup>3</sup> The maximum water holding capacity is calculated with the measurements from soil surveys from the first soil layer by the National Forest Inventory, and generalised to 500m<sup>2</sup> resolution maps (*Laboratoire SILVA, n.d.*)

- edaphic: plant-bioindicated soil pH, C/N ratio, maximum water holding capacity;
- climatic: mean annual temperature and precipitation;
- topographic: elevation, slope, aspect.

Three other environmental composite variables were also extracted or calculated:

- a site index extracted from Toïgo et al., (2015)’s model predicting tree growth as a function of tree species and environmental variables, to account for site productivity;
- the forested region or “*sylvoécorégions*” accounted for the regional context of the plots. It is a classification of forested areas in metropolitan France within which factors discriminant for forestry and habitat distribution are homogeneous (L’IF n°14, 2011) ;
- the plot’s forest type which was created based on an adaptation of an existing French classification, the *BD forêt* version 2 (IGN, 2016a). This classification establishes forest types for plots based on the relative tree cover of each species present on said plot. We approximated tree cover with tree basal area (Figure S1. 4), as tree cover was not accounted for in the reserve inventories. As an example, a plot was considered “pure” when over 75% of the total basal area of the plot was composed of a single species (or a grouping of species with similar characteristics, see Appendix S4 and Figure S1.3 for more details).

The data concerning the nature of the last harvest was not consistent over the plots and only concerned a fraction of the dataset and was not sufficient to build a variable characterising the intensity of harvesting, which is why we settled not to include this information (see Discussion).

### 2.3 Statistical methods

We proceeded to data exploration following (Zuur et al., 2010). All statistical analyses were processed in R version 3.6.3 (R Core Team, 2020). We modelled the time since the last harvesting (dependant variable) as a function of stand structure and environmental variables using a generalized linear mixed model, with gamma error distribution, log link, and a “site” random effect, to take into account the nested sampling design (Bolker et al., 2009). We used the R package *glmmTMB* version 1.0.1 (Brooks et al., 2017) for modelling.

The model aimed at pinpointing maturity features relevant to characterising the time since the last harvesting. We used ascendant variable selection based on AIC (Akaike Information Criterion) scores to select the best model. Considering the large size of our dataset, we chose a conservative five-point AIC threshold instead of the standard two-points threshold. First, we selected stand structure variables, to which we added environmental variables and interactions (see Table 4). We tested first order interactions as well as a few second order interactions that made biological sense. We favoured more specific variables over more generalized ones (e.g. standing and downed deadwood volumes separately vs. total deadwood volume). We checked for correlation and multicollinearity of model covariates, with variance inflation factors (VIF) under five (Zuur et al., 2010).

**Table 4:**
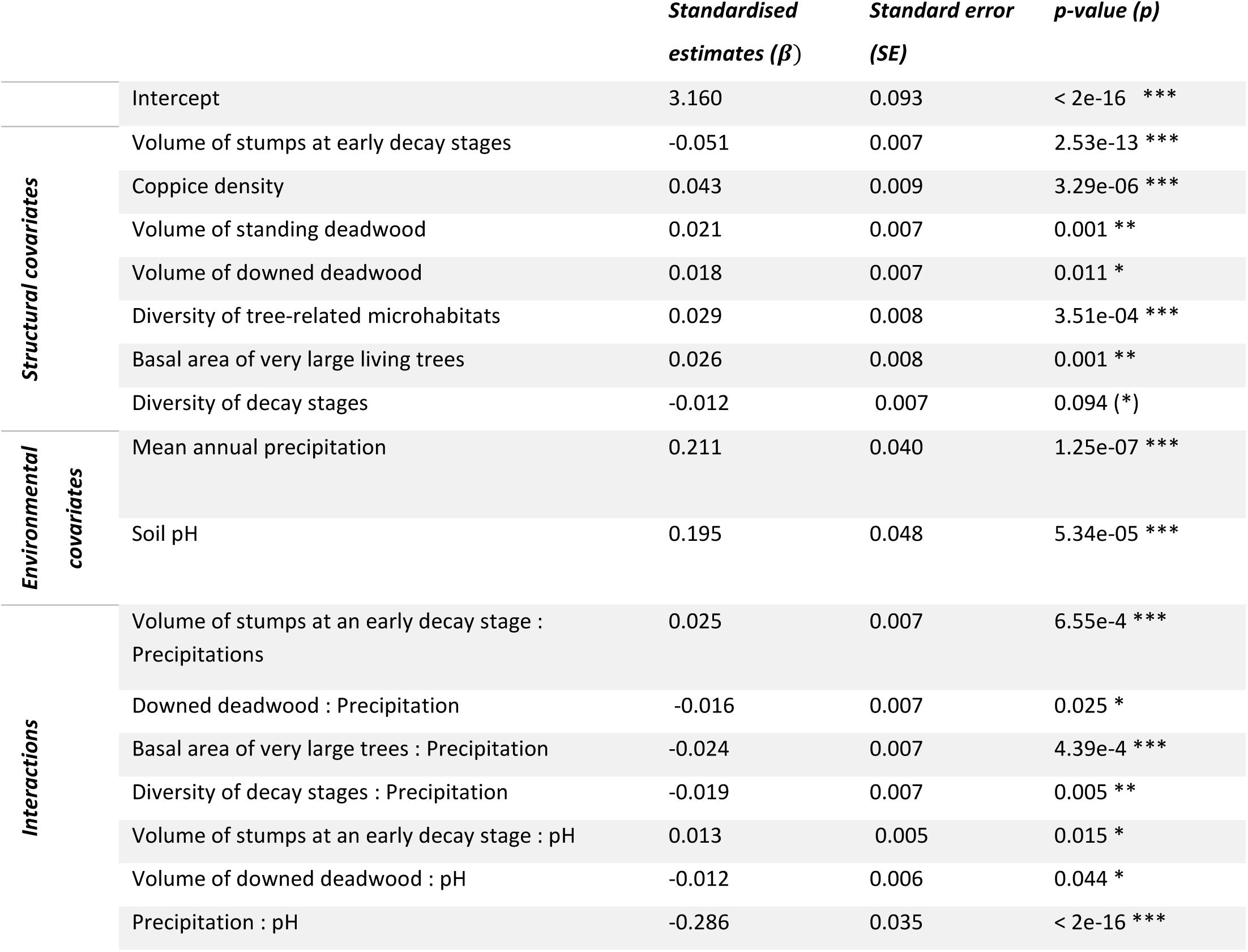
Standardised estimates, standard errors, and p-values for the variables from the generalised linear mixed model with a gamma distribution, log link and “site” random effect explaining the time since the last harvesting of French forest plots. ‘***’ < 0.001, ‘**’ < 0.01, ‘*’ < 0.05, ‘(*)’ <0.1. Volumes and basal areas are all per hectare values.

We validated the final model following (Zuur & Ieno, 2016) by plotting residuals against fitted values, variables included and variables excluded from the model. We checked for overfitting with 50 iterations of 10-fold cross validation (i.e. random sets of 10% of the plots were consecutively removed from the training dataset). Variance explained by the model was evaluated using a pseudo r-squared (Nakagawa & Schielzeth, 2013) with the *piecewiseSEM* package (Lefcheck, 2016). This gives a pseudo r-squared measure for generalised linear mixed model, yielding a marginal r-squared which is the variance explained by the fixed factors, and a conditional r-squared, variance explained by the fixed and random effects. We looked at estimate and prediction accuracy (measure of the coefficient of determination r-squared from the linear regression of predicted versus observed time since the last harvesting) variations (Harrel, 2015). Model robustness was also considered by leaving out reserves one by one and running the model to check for influential or problematic reserves.

### 2.4 Prediction on systematic nation-wide dataset

We predicted the time since the last harvesting on a systematic nation-wide National Forest Inventory (NFI) dataset. Each year, about 6500 NFI forest plots are surveyed, recording measurements of stand structure, soil related and floristic variables. We inferred time since the last harvesting on an NFI subset from the 2012 to 2018 survey campaigns, assuming that most of the plots included have not been subjected to major disturbance during this six-year period, so that our estimations can be valid currently. These plots include both public and private forests, for faithful representation of French forests. We selected 27 075 plots out of the 38 432 NFI plots available, distributed over the whole French territory using the same selection criteria as for training plots (e.g. forest types, see map Fig. S5.1). Since the sampling design of NFI plots is not homogeneous for all regions and forest types, we weighted the time since the last harvesting estimations by the relative proportion of each forest type per forested region (*« sylvoécorégion »*) to account for this heterogeneity.

For the prediction, the final model was simplified as the NFI data was missing some forest attributes included in the initial model fitted on the learning dataset (e.g. tree-related microhabitats, volume of stumps). We ran a new variable selection process with only the variables available in the NFI dataset, which led to the simplified model. Resulting raw model estimates were then used for prediction where we dropped the random site effect since it did not apply to the non-nested NFI data and used a Generalised Linear Model with a gamma distribution and log link. We then calculated the 95% confidence intervals of the raw model estimates using the variance-covariance matrix from the fitted model.

## 3 RESULTS

### 2.1 Description of the training dataset

Our training dataset plots displayed a mean volume of deadwood of 25 m^3^/ha, which is above the national average for similar forest types - about 18 m^3^/ha. The volume of standing deadwood was of 13 m^3^/ha for our plots, against 8 m^3^/ha for the equivalent National Forest Inventory plots. Finally, the mean volume of living trees on our plots was of 303 m^3^/ha, against a national average of 208 m^3^/ha.

Our dataset showed an overall decreasing trend for the number of plots as the time since the last harvesting got longer, and especially few plots with abandonment times over 75 years since the last harvesting (Appendix S1). The median time since the last harvesting was of 26 years.

### 3.2 Selected model

The volume of stumps at an early decay stage was negatively correlated to the time since the last harvesting (standardised β= -0.051, SE = 0.007) (Table 4). The second most important variable, coppice density, had a slightly milder magnitude and was positively correlated to the time since the last harvesting (β= 0.043, SE = 0.009). Other positively correlated variables, but at a lower magnitude than coppices, were the basal area of very large trees (β= 0.026, SE = 0.008), the volume of standing deadwood, i.e. snags and standing dead trees (β= 0.021, SE = 0.007), the volume of downed deadwood (β = 0.018, SE = 0.007) and the diversity of tree-related microhabitats (β = 0.023, SE = 0.008). Furthermore, diversity of decay stages had a negative influence on the time since the last harvesting, but was only marginally significant (β = -0.012, SE= 0.007).

Concerning the environmental covariates, soil pH had a strong positive influence on the time since the last harvesting (β = 0.195, SE = 0.048). The same went for mean annual precipitation (β = 0.211, SE = 0.040). Precipitation and pH were the two effects with the strongest magnitude.

The estimates also showed that when precipitation or pH increased, the negative effect of the volume of stumps on the time since the last harvesting decreased (closer to zero) (β = 0.025, p = 0.007; β = 0.013, SE = 0.005, respectively) and the positive effect of the volume of downed deadwood decreased (β = -0.016, SE = 0.007 ; β = -0.012, SE = 0.006, respectively) (Figure 2). Positive effect of the basal area of very large trees also decreased with higher precipitation (β = -0.024, SE = 0.007). Furthermore, the negative influence of the diversity of decay stages on the time since the last harvesting decreased with more abundant precipitation (β = -0.019, SE = 0.007). Finally, when the soil pH increased, precipitation had a decreasing positive effect on the time since the last harvesting (β = -0.286, SE = 0.035) (see Fig.S5.2 for other illustrations).

**Figure 2:**
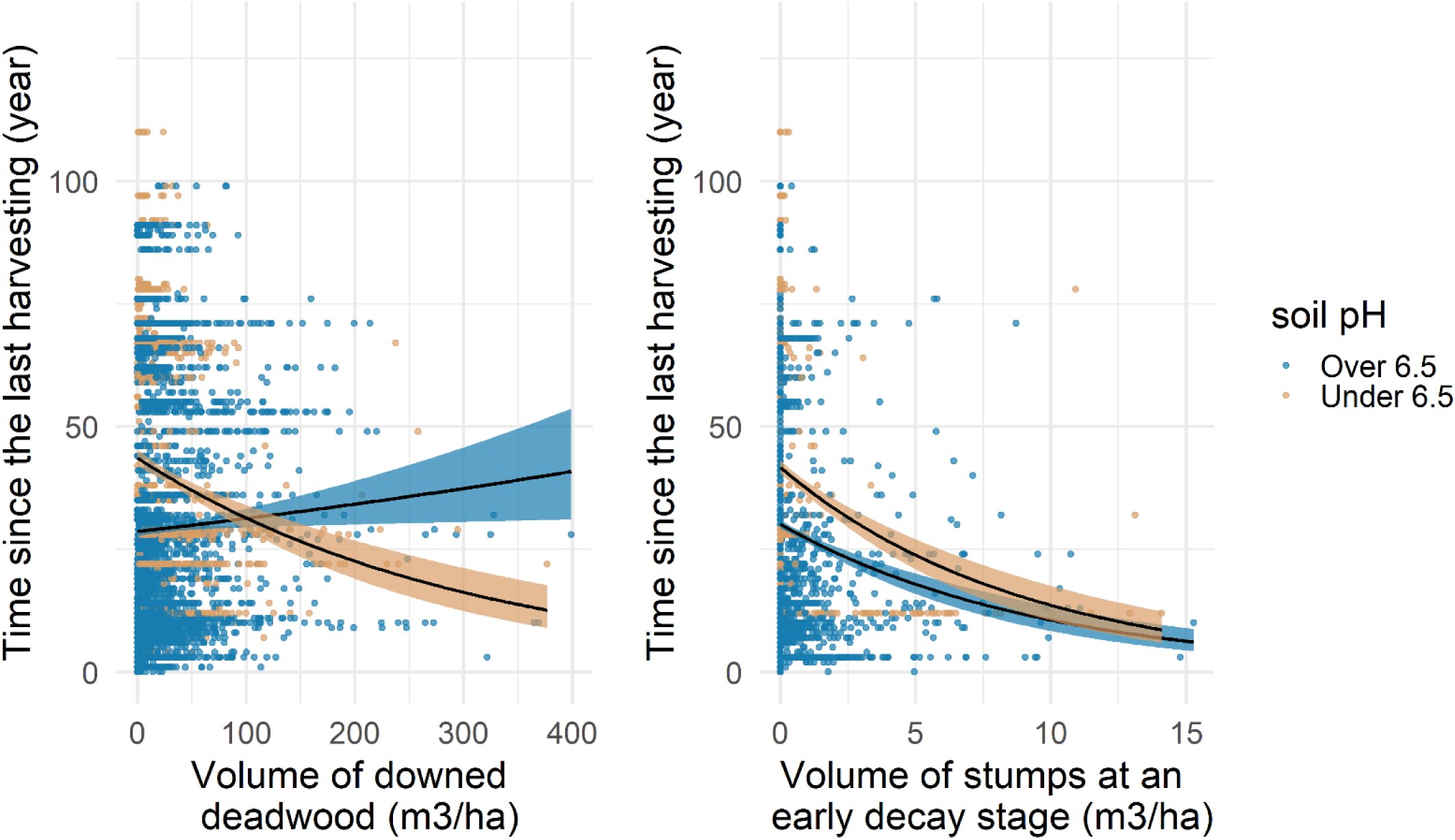
Illustration of the interactions between volume of downed deadwood (β = -0.012, p = 0.044) and volume of stumps at an early decay stage (β = 0.013, p = 0.015) with soil pH. Lines represent a generalised linear model with a log link and 95% confidence interval for pH values over and under 6.5.

### 3.3 Model validation

The distribution of model residuals as a function of the predicted time since the last harvesting showed relative homogeneity, except for a few plots with overestimated time since the last harvesting (Fig. S6. 1). When checked, the residual distribution as a function of the variables both included and excluded from the model showed no noticeable patterns, except for some badly estimated time since the last harvesting on the lower tail of the x axis for the variables “volume of stumps at early decay stages”, “volume of downed deadwood”, “volume of standing deadwood”, and “coppice density”. The pseudo r-squared yielded a marginal r-squared of 0.45 and a conditional r-squared of 0.68.

Reserves were removed one at a time from the training data to check for any particularly influential sites regarding the estimated values. Cross validation results showed that predictions were relatively constant, with an r-squared linking the predicted values to the observed values varying between 0.756 and 0.900 over 500 simulations (Fig. S6.2). The estimated values were quite stable except for the covariates soil pH and precipitation, as well as for the interaction between those two variables for which estimates slightly varied during simulations, but to a negligible extent (Fig. S6. 3).

### 3.4 Model projection on the National Forest Inventory dataset

The simplified model used for projection (see Table S6.1) was comprised of four structural features: the volume of standing and downed deadwood, coppice density and basal area of very large trees, as well as the two environmental covariates soil pH and precipitation. As for interactions, downed deadwood and basal area of very large trees interacted negatively with precipitation, and coppice density interacted negatively with soil pH. Finally, there was still a negative interaction between soil pH and precipitation.

We predicted an average time since the last harvesting of 27 years for the NFI plots issued from the 2012 to 2018 campaigns (including only the six forest types kept for modelling). According to this projection, about 3.1% of French metropolitan forest has reached or surpassed 50 years of abandonment. 43% of forests were predicted to have a time since the last harvesting between 26 and 50 years. When varying this threshold from 30 to 300 years of abandonment, we observe the decreasing trend presented in Figure 3a. The confidence interval is very large, around 30 years without harvesting and gets narrower with higher thresholds. There are very few forests predicted above 75 years without harvesting, which is in accordance with the distribution of the time since the last harvesting in the training dataset (French forest reserves). Plots with longer predicted time since the last harvesting are mainly distributed in mountainous regions of eastern France (Vosges, southern Alps and Jura), as well as in the south of France (Pyrenees and Cevennes) and Corsica (Figure 3d).

**Figure 3:**
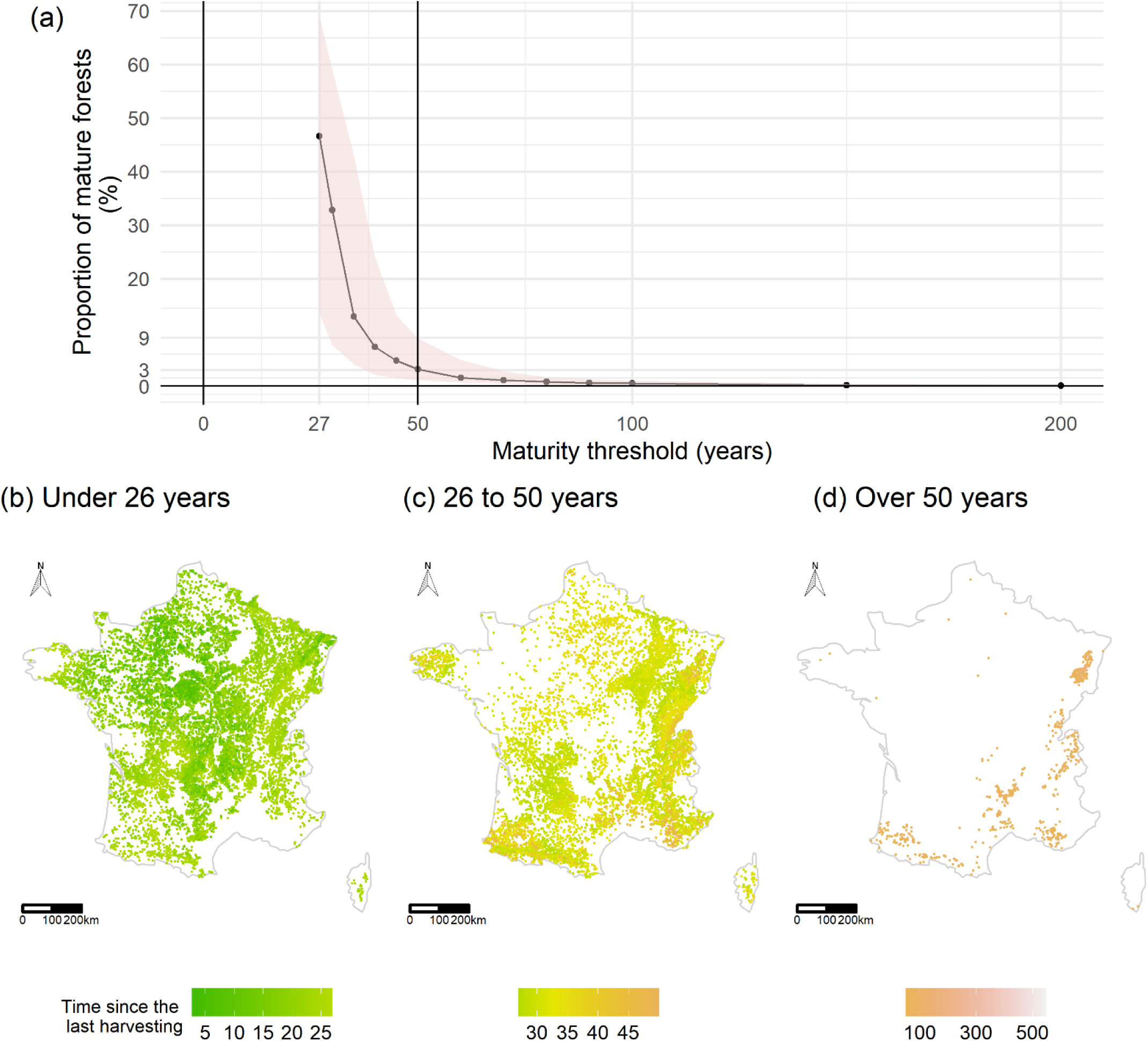
(a) represents how the maturity threshold changes the predicted proportion of overmature forests. The ribbon represents the 95% confidence interval. The 50-year threshold (vertical bar) is the one we chose for this study. The confidence interval narrows very fast and at 50 years without harvesting it ranges from 1 to 9%. (b) NFI plots with predicted time since the last harvesting under 26 years, (c) NFI plots with predicted time since the last harvesting between 26 and 50 years, (d) NFI plots with predicted time since the last harvesting over 50 years.

## 4 DISCUSSION

We evaluated French forest’s state of maturity using time since the last harvesting and showed that most maturity features highlighted in the literature did indeed explain time since the last harvesting quite well, and that density of coppices, translating legacy of past forest management, also played a role in the modelling approach. Our projection gives the first robust statistical estimate of the proportion of overmature forests in metropolitan France and may serve to report on their status. Our approach allowed us to account for the multidimensional and continuous nature of forest maturity, and differs from previous studies (e.g. Larrieu et al., 2019a; Paillet et al., 2015; Vandekerkhove, 2005) which used the opposite reasoning: modelling structural variables as a function of the time since the last harvesting.

### 4.1 Influence of parameters on the models

As expected, the volume of stumps at early decay stages was negatively correlated with the time since the last harvesting (Paillet et al., 2015; Siitonen et al., 2000). When plots are unharvested for a long period of time, stumps that originated from human disturbance decompose, hence the decline in abundance of fresh stumps in more overmature forests. Stumps at an early decay stage are characteristic of recent harvesting, and their absence is a token of longer abandonment times.

Higher basal area of very large living trees is also a known characteristic of overmature forests (Bauhus et al., 2009; Burrascano et al., 2013; Paillet et al., 2015). Additionally, larger trees often bear higher diversity of tree-related microhabitats and therefore promote biodiversity at the stand level (Larrieu et al., 2018; Paillet et al., 2017): tree-related microhabitats are the result of tree alteration by biotic and abiotic processes (Larrieu et al., 2018), their presence is thus mostly related to the timespan during which trees are subjected to those processes and thus to time since the last harvesting.

We also expected the diversity of decay stages on standing and downed deadwood to be positively correlated to the time since the last harvesting. Although it was, the observed effect was only marginally significant. Wood decomposition dynamics depend on other factors than time only, such as environmental conditions prone to microorganisms (Herrmann & Bauhus, 2013), the chemical composition of wood (i.e. species), its diameter, micro-climatic conditions as well as edaphic parameters (Heiri et al., 2009; Přívětivý et al., 2018; Larrieu et al., 2019b). Those characteristics were not selected in the model interactions, it therefore seems that there is no univocal correlation between the time since the last harvesting and decay stages, hence the marginal effect observed here. Moreover, secondary disturbances generating deadwood may vary in nature, severity and temporality, meaning that even in overmature forests, deadwood characteristics can be variable (Brassard & Chen, 2006). Some punctual disturbances such as storms or droughts can result in locally high amounts of deadwood (Aakala, 2011; Harmon, 2009). In our case, some plots have been subjected to major disturbances, e.g. the 1999 storms that caused damage to a large portion of the French forests, which could still bear the legacy of those events. Additionally, managed stands are usually harvested decades before they reach “true” old-growth stage, meaning that recently abandoned forests will hold deadwood at relatively low decay stages and may need several decades to have an important diversity of decay stages. Thus, Wirth et al. (2009) suggested that deadwood volume and decay stages could potentially be misleading and could not be taken as sole indicators of maturity. Nevertheless, our results confirm that deadwood is a key component for characterising overmature French forests, but that they may still be at too early successional stages to see the true shape of the relationship between diversity of decay stages and the time since the last harvesting. This also might be more complex than anticipated considering the diversity of abiotic and biotic processes that come into play in deadwood decay rates and accumulation (Harmon, 2009).

Finally, coppice density was strongly and positively correlated to the time since the last harvesting. Coppicing is a legacy of past management for fuel (heating) and is a practice that has decreased over the last century (IGN, 2016b). Many coppice-with-standards plots have now been abandoned (Unrau et al., 2018) resulting in what Lassauce et al. (2012) call “coppice-with-standard stands with an overmature component”. In their study, they also found a significant increase in the basal area of large living trees and downed deadwood for more overmature plots with a history of coppicing. This trend seems widespread among stands with a history of coppice-with-standard management. Indeed, Becker et al., (2017) also noticed that after 40 years of abandonment of coppicing, legacies of this practice still remained in species composition. These overmature coppices could constitute another form of overmature forests (Lassauce et al., 2012), and it would be interesting to consider coppice density as another indicator for forest maturity, especially since coppice forests currently represent up to 15% of Europe’s forest resources (Unrau et al., 2018).

Environmental variables were amongst the strongest positive predictors for the time since the last harvesting. We related the importance of mean annual precipitation and soil pH to soil fertility : soil nutrients need to be in a soluble form and are most available at a neutral soil pH of 6.5 to 7.5 (Jense, 2010; Landsberg & Gower, 1997). The forest reserve plots have a mean soil pH of 5.3, meaning that increased pH in our plots corresponds to optimum nutrient absorption potential, hence the observed positive correlation between soil pH and time since the last harvesting. Surprisingly, elevation was not selected in favour of mean annual precipitation). Precipitation was nonetheless strongly correlated to elevation (correlation coefficient = 0.9). Precipitation seems to have a broader role and a higher explanatory power than elevation, since they condition both biotic (tree growth, decay rates, soil properties) and abiotic (atmospheric humidity) phenomena. Indeed, environmental variables interacted significantly with stand characteristics in the model, notably interactions between deadwood volumes and pH/precipitation. Plots with higher pH and higher precipitation (better site productivity), have the potential for high basal areas and high stem densities to be reached sooner. Nonetheless, we observed a negative interaction between soil pH and precipitation, meaning that overall, productive sites show shorter times since the last harvesting. Rainier plots and/or plots with higher soil pH conditions showed lessened negative effects of the volume of stumps, and positive effect of the diversity of decay stages, the volume of downed deadwood and basal area of very large trees on the time since the last harvesting. These interactions could be linked to either (1) known productive sites (higher precipitation and soil pH) being more attractive for harvesting, e.g. productive sites with high increment and larger trees, which would be favoured for harvesting, or (2) sites with heavier rainfall (> 1000 mm) are subjected to faster decomposition rates of deadwood (Zell et al., 2009) compared to drier surroundings, resulting in misleadingly low time since the last harvesting for those rainier plots.

### 4.2 The projected proportion and rough distribution of overmature forests

Our model predicted that 3.1% of the national forested area has a time since the last harvesting above 50 years (95% confidence interval ranges from 1 to 9 %). This prediction is valid only for the six forest types kept for modelling (pure beech, pure deciduous oak, mixed broadleaved, pure spruce and/or fir, mostly broadleaved and mostly coniferous types). The mean prediction is of the same order of magnitude as the early expertise that had been proposed in 1993 (MAAPRAT-IFN, 2011). The fact that this proportion has not changed since the 90s is somewhat surprising but given that this former estimation came with no statistical background, it is quite an unreliable figure to compare to. The only recent reliable numbers are those reported concerning the area of primary forests, which account for less than 0.1% of the national forested area (Barredo et al., 2021; Sabatini, Bluhm, et al., 2020).

As expected, forested areas abandoned more than 50 years ago seem to occur mostly in more remote and mountainous areas in southern and eastern France (Figure 3c). Indeed, it has been shown that accessibility and favourable topographic, climatic, and soil conditions characterise intensively managed areas, whereas de-intensification and abandonment trends occur in more marginal areas (Levers et al., 2014, 2018; Sabatini et al., 2018) Additionally, abandoned forests have been found to be located in low productivity and low accessibility areas in the past (Joppa & Pfaff, 2009; Lõhmus et al., 2004; Svensson et al., 2020).

More interestingly, our prediction also showed evidence of a large proportion (43%) of French forest with abandonment times comprised between 26 and 50 years, with “hotspots” such as Corsica, Brittany and middle-eastern France. This figure is however to be tempered, as it comes with considerable uncertainty, the lower bound of the confidence interval being around 12.5%. For example, the spatial pattern in the projection of time since the last harvesting in the area of Britany (West of France) could be linked to the scarcity of training data for this area combined with the fact that it is a less forested region with high precipitation, making the model particularly sensitive and probably not quite fitted for this particular region.

Nonetheless, they represent a significant proportion of French forests and could - in a relatively close future - display interesting conservation attributes which would deserve more attention, especially if some larger continuous and connected areas, particularly interesting for biodiversity, could be restored or managed sensibly (Bauhus et al., 2009; Portier et al., 2020; Sabatini et al., 2020). We mainly associate this large proportion with the fact that French metropolitan forested surface area has doubled over the last century mostly due to land abandonment and now occupies 31% of the metropolitan France or about 16.9 million hectares (IGN, 2018). While strict forest reserves compose a mere 0.15% of the French forests (Cateau et al., 2017), private forests represent 75% of the area (MAAF & IGN, 2016) and considering that the harvesting in these - often small and fragmented - properties may be low or inexistent, they could potentially account for this large portion of “in between” forests in our prediction. In addition, about 60% of the annual biomass increment is actually harvested in France, a phenomenon that feeds the proportion of forests unharvested for an intermediate timespan. Finally, many of these “in between” forests are in mountainous areas known for their potential for matured forests (see above).

It is interesting to note those “in between” areas which are not accounted for in most studies and conservation decisions. If we consider that each country is equally responsible for contributing to the “EU Biodiversity Strategy for 2030” goal for 10% of strictly protected land (European Commission, 2020) and we project this figure on the forest ecosystem, then France, with 3% of estimated overmature forest – not all protected and probably quite fragmented - still has a long way to go. But the recognition that areas with high potential for achieving this goal exist and are identified, is a first step forward.

### 4.3 Limits and perspectives to our modelling approach

The distribution of overmature forests predicted by our model should not be taken as a precise map but as an overview of the state of French forests to this day. Indeed, the plot density is quite scarce with 27 075 plots for the whole French territory. The time since the last harvesting used for training the model could introduce some error, since the older the last known harvesting, the more uncertain this estimation becomes. Although the integration of the volume of coppice in our model is a way to take into account at least part of the harvesting legacy of the plot, we acknowledge that our model does not consider the intensity of harvesting (e.g. volume of wood harvested, type of management), and – had this knowledge been available – it would likely have further refined our predictions. In this regard, private forests issued from land abandonment would differ from forests already in place with similar time since the last harvesting. One way to take this into account without knowing the plot history would be to offset our prediction with maps of ancient forests (forests in place since the 1850s), the digitization of which is currently a work in progress (Bergès & Dupouey, 2021). Another consequence is related to the possible interaction between the silvicultural treatment and tree related microhabitats. Some trees bearing microhabitats are systematically removed during thinning operations while stands managed as coppice-with-standards often have smaller trees, that could be too small to bear many tree-related microhabitat (Paillet et al., 2017).

Additionally, our model would have been more precise had we not removed covariates from the predictive. This is further proof of the use of surveying stumps and tree-related microhabitats in forest inventories (Paillet et al., 2017).

The overall results of our study are encouraging, in the sense that a large portion of the French forest has potential to attain interesting states of maturity in a close future and therefore to contribute to the race against climate change, by acting as carbon pools (Carey et al., 2001; Portier et al., 2020; Sabatini et al., 2020). In some cases high carbon storage may even support conservation of biodiversity though trade-offs between carbon stock and biodiversity have also been found (Sabatini et al., 2019). Further validation, e.g. by crossing our findings with maps and data on previously studied overmature sites, along with field validation, would enable us to refine our knowledge of the distribution of more overmature forests, and evaluate the precision of our predictions. These overmature forests complement the integrated conservation measures in managed forests and constitute a functional network for forest biodiversity, the efficiency and completeness of which remains to be analysed (Vandekerkhove et al., 2013).

Our results can be used in ensuring that the existing 3% of overmature forest are acknowledged, and our work can constitute a stepping-stone to further refining our knowledge of the state of maturity of French forests. On a broader scale, similar methods could be applied for neighbouring temperate North-western European forests with characteristics and harvesting history alike that of French forests (Sabatini, Bluhm, et al., 2020; Sabatini et al., 2018). Indeed, since many countries around the world benefit from national forest inventories that provide robust forest estimates and could be used as independent data to project results of models such as ours (see Tomppo et al., 2010 for a synthesis). The limiting factor is probably the fact that time since last harvesting is not often documented at a large scale or requires deep historical work to be gathered. We assume that our model could roughly be applied to neighbouring countries with similar ecological conditions and forest types (e.g. Germany, Switzerland). Beyond, it should be necessary to recalibrate the model with sound and local independent field data. Would this process be applied at a large scale, it would be a complementary source of knowledge to strengthen estimates from other initiatives (Sabatini et al., 2018; Sabatini, Keeton, et al., 2020) and thus provide more decision tools for the conservation of primeval and overmature forests that have a crucial role for biodiversity and mitigation of climate change (Luyssaert et al., 2008; Paillet et al., 2010).

## 5 ACKNOWLEDGEMENTS

This study is issued of the Master’s training period of LT and was funded by the French Foundation for Biodiversity Research (“Appel Masters 2019”, www.fondationbiodiversite.fr) and the Direction for Water and Biodiversity of the French ministry in charge of the environment (convention “FORETMAT”). We are in debt to the reserve and forest managers who fed the databases and made this study possible. Without their daily commitment and implication in the forest surveys, we would not have reached such complete and global results. We are also grateful to F. Benest (IGN) and B. Courbaud (INRAE) for constructive discussions on the different parts of this work. Finally, we thank the reviewers for their invaluable addition to our work.

## 6 DATA ACCESSIBILITY

The two datasets we worked with for this study were the Nature Forest Reserve data (PSDRF), ONF-RNF and the French national forest inventory data (raw data, annual surveys from 2005 to 2018, https://inventaireforestier.ign.fr/spip.php?rubrique159).

