## Supplementary material for "How much does it take to be old? Modelling the time since the last harvesting to infer the distribution of overmature forests in France"

### SUPPLEMENTARY MATERIALS

### Thompson et al. 2021

Appendix S1:

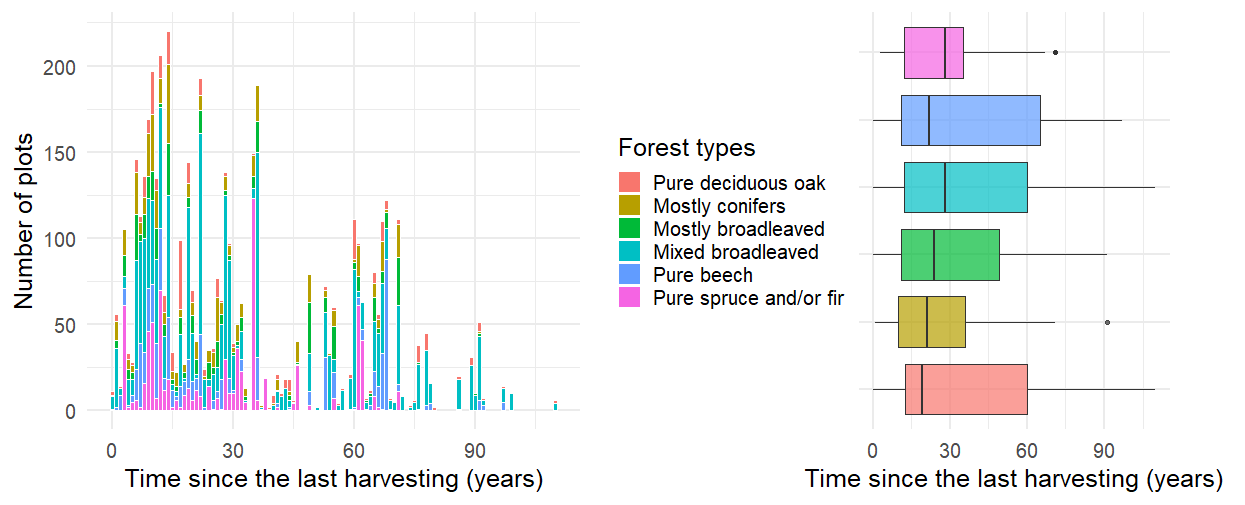

Figure S1. 1: Distribution of time since the last harvesting of the reserve database plots coloured by forest type.

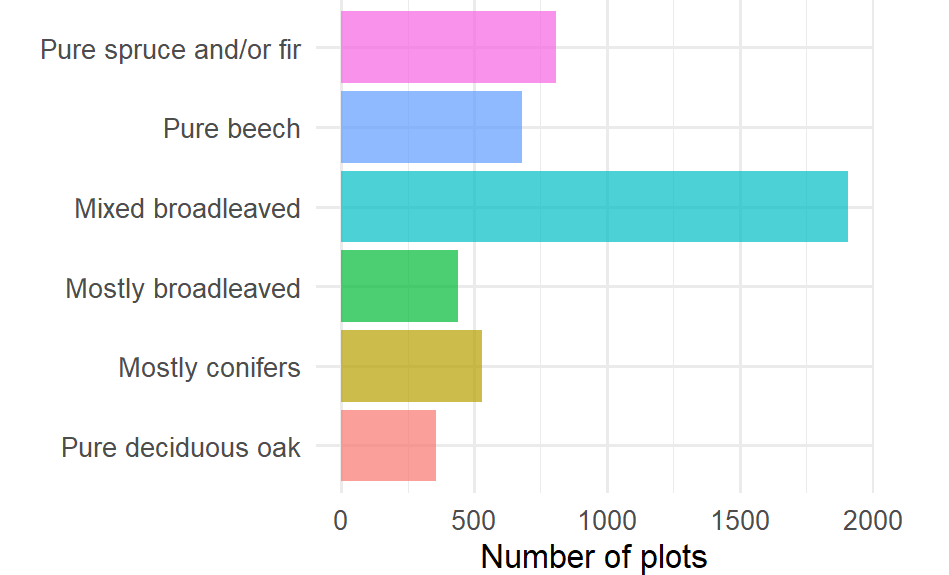

Figure S1. 2: Six most abundant forest types among the 4728 ploys kept from the PSDRF data. “Pure » types: species group represents over 75% of total basal area; « Mostly » types: > 50% of total basal area; « mixed » types: > 75% of total basal area.

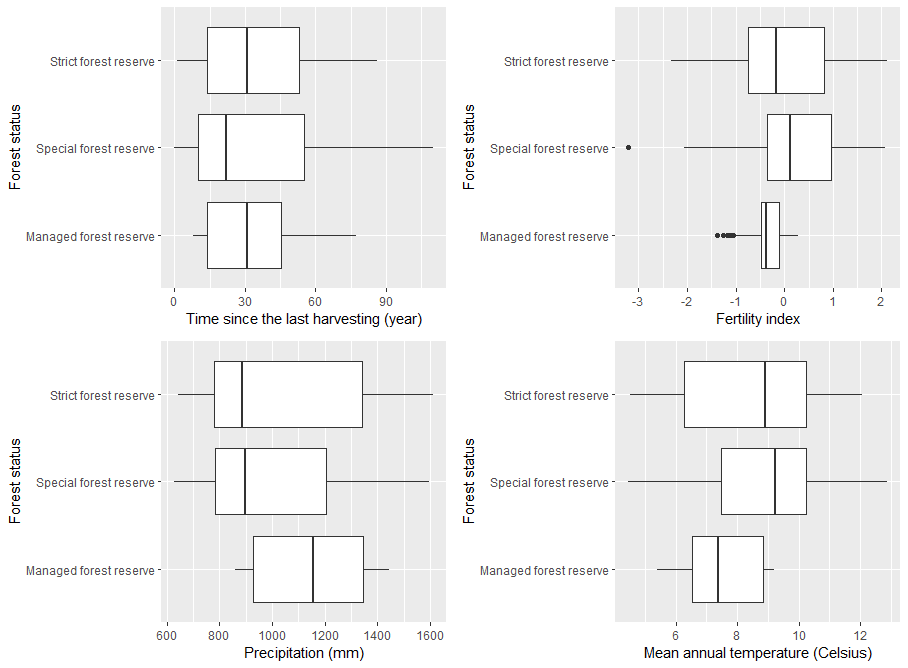

Figure S1. 3 : Comparison of site fertility, temperature, precipitation grouped by management regime

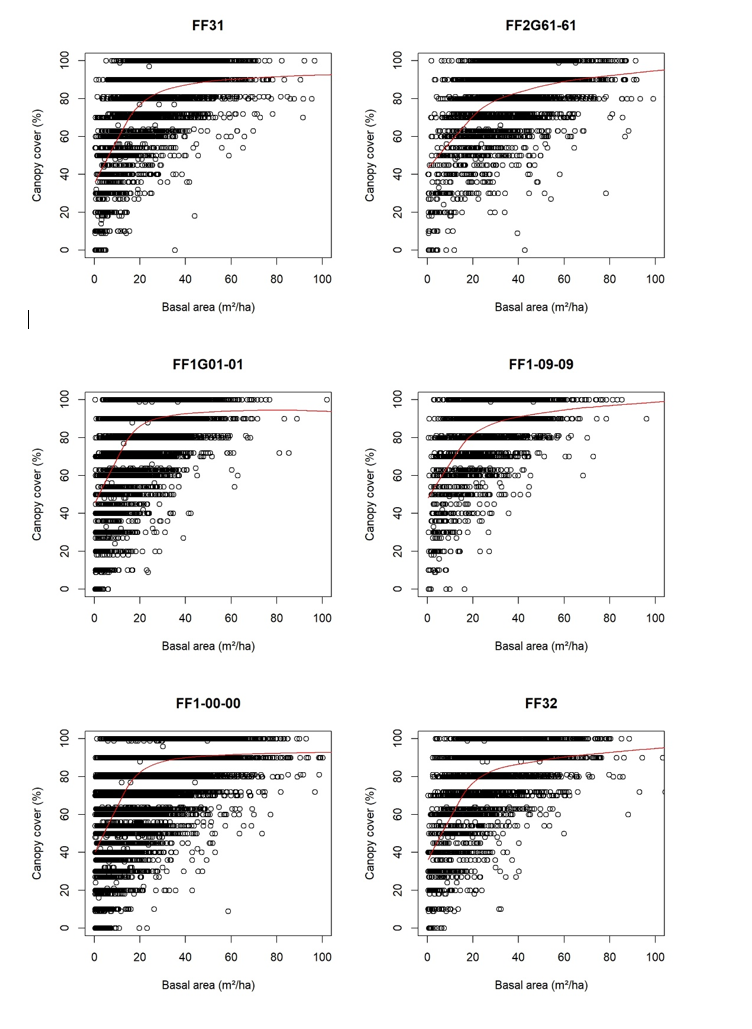

Figure S1. 4 : Relationship between tree basal area and canopy cover for the 6 forest types (See Appendix S3 for correspondence between code and forest type)

*Appendix S2: Protocol for the French nature forest reserves stand structure description*

On each circular plot, forest stand structure was characterized using the following protocol (Paillet et al., 2015, 2019):

- All living trees larger than 7.5 cm were measured considering, by checking if they issued from the same base or not, the difference between coppice and standard trees.
- For living trees with a diameter at breast height (DBH) above 30 cm, a fixed angle plot method was used to select the individuals within a relascopic angle of 3%. Practically, this meant that sampling distance was proportional to the apparent DBH of a tree. For example, a tree with a DBH of 60 cm was included in the sample if it was within 20 m of the center of the plot. This technique allowed us to better account for larger trees at a small scale.
- All other variables were measured on fixed-area plots. Within a fixed 10-m (314 m2) radius plot, the diameter of all living trees and standing deadwood (stumps, snags (height > 130 cm), standing dead trees) was measured from 7.5 to 30 cm DBH.
- Within a 20-m radius (1256 m2), all standing deadwood and downed deadwood (logs) with a diameter > 30 cm were recorded.
- Downed deadwood (logs) with a diameter under 30 cm was recorded along three separate 20 m long radial transects (every 127°).
- All standing trees (alive and dead) were visually inspected for the recording of tree-related microhabitats. Observers attended a training session and were given a field guide with pictures to help them better determine microhabitat types and detailed criteria to include in the inventories.

On every piece of deadwood decay stage of the bark and of the wood were recorded, respectively on a scale of one to four and one to five. Whenever possible, all trees, both alive and dead, were identified to species level.

**Decay stages determination for deadwood**

Bark decomposition is assessed visually :

1: bark on the whole piece

2: bark covering more than 50% of the piece

3: bark covering less than 50% of the piece

4: no more bark

Decay stage is assessed with knife-test :

1: Hard or non-altered

2: < ¼ of the diameter decayed

3: ¼ to ½ of the diameter decayed

4: ½ to ¾	of the diameter decayed

5: > ¾ of the diameter decayed

Appendix S3: Table presenting the classification used to derive plot the harmonized tree related microhabitat classification. The first column is the label from (Larrieu et al., 2018)’s types simplified into their “forms” which we used to regroup our four classifications. Ø = diameter. NA stands for the forms that were not inventoried by the different protocols.

| Form | Type | Threshold |
| --- | --- | --- |
| Breed.Woodpecker | Small woodpecker breeding cavity (ø < 4 cm) | Cavity entrance ø < 4 cm |
| Breed.Woodpecker | Medium-sized woodpecker breeding cavity (ø = 4-7 cm) | Cavity entrance ø = 4-7 cm |
| Breed.Woodpecker | Large woodpecker breeding cavity (ø > 10 cm) | Cavity entrance ø > 10 cm |
| Breed.Woodpecker | Woodpecker “flute” (string of ? 3 breeding cavities ) | Cavity entrance ø > 3 cm |
| Rot.Hole | Trunk base rot hole | Opening ø >10 cm |
| Rot.Hole | Trunk rot hole | Opening ø >10 cm |
| Rot.Hole | Semi-open trunk rot hole | Opening ø > 30 cm |
| Rot.Hole | Chimney trunk base rot hole | Opening ø > 30 cm |
| Rot.Hole | Chimney trunk rot hole | Opening ø > 30 cm |
| NA | Hollow branch | Opening ø > 10 cm |
| NA | Insect galleries and bore holes | Hole ø > 2 cm or numerous smaller holes covering an area > 300 cm² (A5 format) |
| Concavities | Dendrotelm (phytotelmata, water-filled hole) | ø > 15 cm |
| Concavities | Woodpecker foraging excavation | Depth > 10cm, ø > 10cm |
| Concavities | Trunk bark-lined concavity | Depth > 10cm, ø > 10cm |
| Concavities | Root buttress concavity | Entrance ø > 10 cm |
| Bark.Loss | Bark loss | Area > 300 cm² (A5 format) |
| Bark.Loss | Fire scar | Area > 600 cm² (A4 format) |
| Bark.Loss | Bark shelter | Gap > 1 cm depth > 10 cm height > 10 cm |
| Bark.Loss | Bark pocket | Gap > 1 cm width > 10 cm height > 10 cm |
| Exposed.HeartWood | Stem breakage | Stem ø > 20 cm at the broken point |
| Exposed.HeartWood | Limb breakage (trunk heartwood exposed) | Area of exposed heartwood > 300 cm² (A5 format) |
| Crack | Crack | Length > 30 cm width > 1 cm depth > 10 cm |
| Crack | Lightning scar | Length > 30 cm width > 1 cm depth > 10 cm |
| Crack | Fork split at the intersection | Length > 30cm |
| Crown.DeadWood | Dead branches | Branch ø > 10cm, or branches ø > 3cm and > 10% of the crown is dead |
| Crown.DeadWood | Dead top | ø > 10cm at the lower part of the piece of deadwood |
| Crown.DeadWood | Remaining broken limb | Limb ø > 20cm at the broken end length of the remaining piece > 0,5m |
| Twig.tangles | Witch broom | Largest ø > 50cm |
| Twig.tangles | Epicormic shoots | > 5 twig clusters |
| Burr.Canker | Burr | Largest ø > 20cm |
| Burr.Canker | Canker | Largest ø > 20cm or large part of the trunk covered |
| Polypore | Perennial polypore | Largest ø > 5cm |
| Polypore | Annual polypore | Largest ø > 5cm or cluster of > 10 fruiting bodies |
| NA | Pulpy agaric | Largest ø > 5cm or cluster of > 10 fruiting bodies |
| NA | Pyrenomycete | Stroma ø > 3cm or stroma cluster covering > 100cm² |
| NA | Slime mold | Largest ø > 5cm |
| Epiphytes | Bryophyte (moss or liverwort) | > 10% of the trunk area covered |
| Epiphytes | Foliose or fruticose lichens | > 10% of the trunk area covered |
| Epiphytes | Ivy or liana | > 10% of the trunk area covered |
| NA | Ferns | > 5 fronds |
| NA | Mistletoe | Largest ø > 20cm for Viscum spp. and Loranthus europaeus more than 10 clusters for Arceuthobium oxycedri |
| Nest | Vertebrate nest | ø > 10cm |
| Nest | Invertebrate nest | Presence (observation of nest or associated insects) |
| Microsoil | Bark microsoil | Presence (direct observation or specific fungi) |
| Microsoil | Crown microsoil | Presence |
| Sap.Run | Sap run | Length > 10 cm |
| Sap.Run | Heavy resinosis | Length > 10 cm |
| NA | Snag base coarse woody debris | Presence (direct observation or specific fungi) |
| NA | Rootplate | Height > 1m |

Appendix S4: Table presenting the classification used to derive plot forest types from plot species composition. This classification is inspired from the “BD forêt version 2” classification, which is based on the species’ relative tree cover. We chose to use tree basal area instead, as tree cover was not unknown in our dataset. Each plot forest type is defined as the proportion of a specific species’ basal area on the said plot. Therefore, plots are called “pure” if over 75% of the total plot basal area is occupied by one (or two in a few cases) species. We then called mixed any plots where a mixture of two or more species occupied over 75% of the total basal area. Finally, the last two broader forest types correspond to plots where a true mixture of tree species occur and the only definition we can give is if either broadleaved or coniferous trees are preponderant. In the end, only the six highlighted types out of the 22 included enough plots to be kept for modelling. The six forest types kept for modelling are: pure deciduous oak, mostly coniferous trees, mostly broadleaved trees, mixed broadleaved trees, pure beech trees, Pure spruce and/or fir.

| Code *BD forêt version 2* | Class | Label | Definition | Species |
| --- | --- | --- | --- | --- |
| FF1G01-01 | Broadleaved | Pure deciduous oak | Basal area of deciduous oaks ≥ 75% | *Quercus cerris, Quercus rosacea, Quercus robur, Quercus pubescens, Quercus rubra, Quercus petraea, Quercus tauza* |
| FF1G06-06 | Broadleaved | Pure evergreen oak | Basal area of evergreen oaks ≥ 75% | *Quercus suber, Quercus ilex* |
| FF1-09-09 | Broadleaved | Pure beech | Basal area of beech trees ≥ 75% | *Fagus sylvatica* |
| FF1-10-10 | Broadleaved | Pure chestnut | Basal area of chestnut trees ≥ 75% | *Castanea sativa* |
| FF1-14-14 | Broadleaved | Pure black locust | Basal area of black locust trees ≥ 75% | *Robinia pseudoacacia* |
| FF1-49-49 | Broadleaved | Other pure broadleaved trees | Basal area of a single other broadleaved species ≥ 75% | Ash tree (*Fraxinus sp*),  Maple trees (*Acer sp*),  Birch (*Betula sp*),  Charm (*Carpinus betulus*),  Alder (*Alnus sp*),  Hazel tree (*Corylus sp*) |
| FF1-00-00 | Broadleaved | Mixed broadleaved trees | Combined basal area of at least two broadleaved species that are not either deciduous oak or evergreen oak ≥ 75% (without any single species occupying a basal area ≥ 75%) | *Fagus sylvatica, Castanea sativa, Robinia pseudoacacia*,  Ash trees (*Fraxinus sp*),  Maple trees (*Acer sp*),  Birch (*Betula sp*),  Charm (*Carpinus betulus*),  Alder (*Alnus sp*),  Hazel tree (*Corylus sp*) |
| FF2-51-51 | Coniferous | Pure maritime pine | Basal area of maritime pine ≥ 75% | *Pinus pinaster* |
| FF2-52-52 | Coniferous | Pure scots pine | Basal area of scots pine ≥ 75% | *Pinus sylvestris* |
| FF2G53-53 | Coniferous | Pure laricio pine and/or black pine | Basal area of laricio pine and/or black pine ≥ 75% (can be either a mixture of both species or each species individually) | *Pinus nigra* |
| FF2-57-57 | Coniferous | Pure Aleppo pine | Basal area of Aleppo pine ≥ 75% | *Pinus halepensis* |
| FF2G58-58 | Coniferous | Pure mountain pine and/or swiss pine | Basal area of laricio pine and/or black pine ≥ 75% (can be either a mixture of both species or each species individually) | *Pinus mugo, Pinus cembra* |
| FF2-81-81 | Coniferous | Other pure pine | Basal area of another pine species ≥ 75% | *Pinus mugo, Pinus pinea* |
| FF2-80-80 | Coniferous | Mixed pine | Combined basal area of at least two pine species ≥ 75% (without being a mixture of black pine and laricio pine or of mountain pine and swiss pine and without any single species occupying a basal area ≥ 75%) | *Pinus pinaster, Pinus sylvestris, Pinus nigra, Pinus halepensis, Pinus mugo, Pinus cembra, Pinus mugo, Pinus pinea* |
| FF2G61-61 | Coniferous | Pure spruce and/or fir | Basal area of spruce and/or fir ≥ 75% (can be either a mixture of both species or each species individually) | *Abies cephalonica, Abies nordmanniana, Abies grandis, Abies alba,*  *Picea abies, Picea sitchensis* |
| FF2-63-63 | Coniferous | Pure larch trees | Basal area of larch trees ≥ 75% | *Larix decidua* |
| FF2-64-64 | Coniferous | Pure douglas fir | Basal area of Douglas firs ≥ 75% | *Pseudotsuga menziesii* |
| FF2-91-91 | Coniferous | Other pure coniferous trees | Basal area of a single other coniferous species ≥ 75% | Cedar trees *(Cedrus atlantica, Cedrus libani, Thuja sp)*  Cypress *(Cupressus sp)*  Yew *(Taxus baccata)* |
| FF2-90-90 | Coniferous | Other mixed coniferous trees | Basal area of a mixture of at least two coniferous species which are not either a mixture of pine trees or spruce and fir ≥ 75% (without any single species occupying a basal area ≥ 75%) | Spruce, fir*,* cedar, cypress, yew (see above) |
| FF2-00-00 | Coniferous | Mixed pine and other coniferous trees | Basal area of a mixture of at least a pine tree and another coniferous species ≥ 75% (without any single species occupying a basal area ≥ 75%) | *All coniferous species above* |
| FF31 | Broadleaved/Coniferous | Mostly broadleaved trees (mixture of broadleaved and coniferous trees) | 50% ≤ Basal area of broadleaved trees ≤ 75% | All broadleaved and coniferous species above |
| FF32 | Broadleaved/Coniferous | Mostly coniferous trees (mixture of broadleaved and coniferous trees) | 50% ≤ Basal area of coniferous trees ≤ 75% | All broadleaved and coniferous species above |

Appendix S5: maps and graphs of the two datasets

Fig. S5.1: Map of the 27 075 National Forest Inventory plots from the 2012 to 2018 campaigns in metropolitan France which served as the spatial prediction dataset. Plots are coloured by forest type.

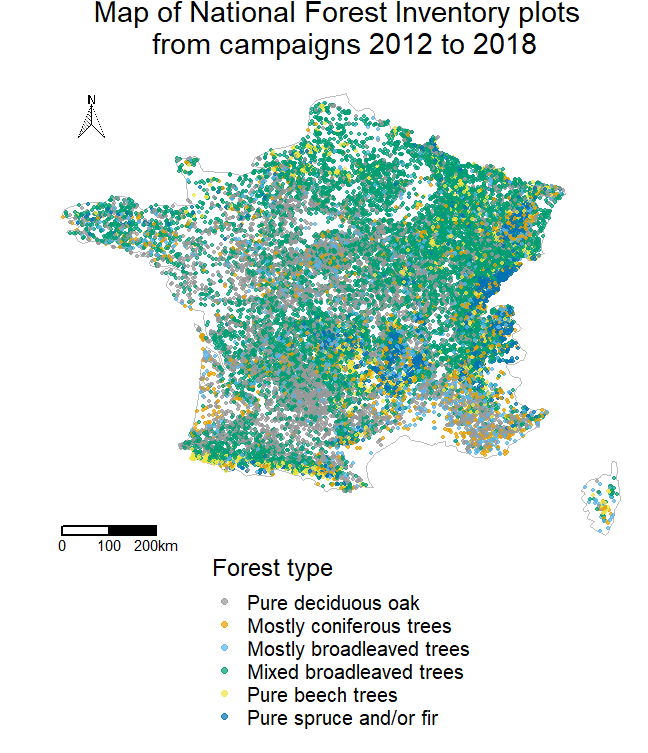

Fig. S5.2: Illustration of the interactions between the maturity features and precipitation level (threshold of 1000 mm is the mean precipitation for the forest reserve plots)

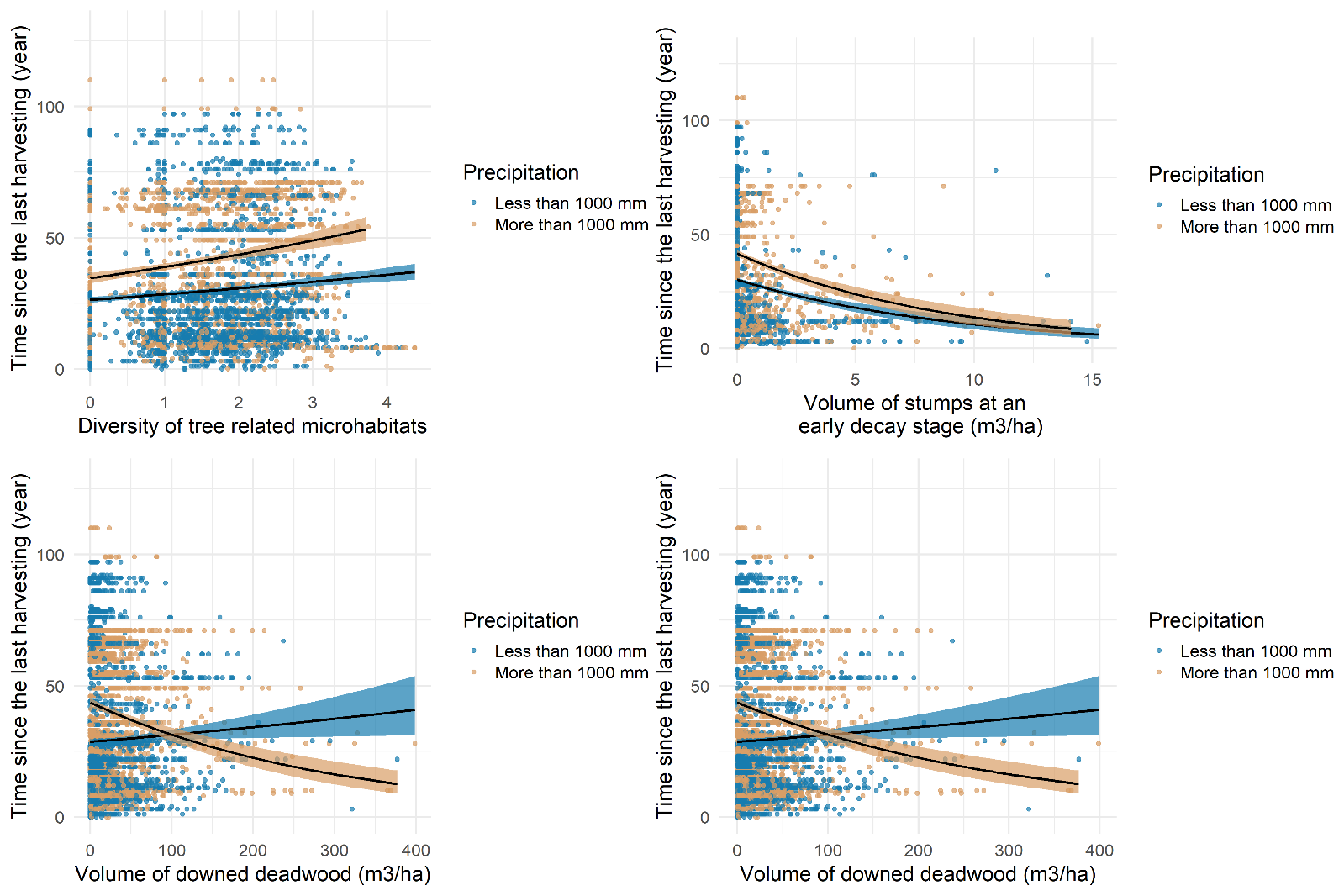

Appendix S 6: model validation

Fig. S6. 1: Distribution of the Pearson residuals from the generalised linear mixed model explaining the time since the last harvesting using structural features and environmental variables. The trend is satisfyingly neutral, although a few plots have overestimated time since the last harvesting.

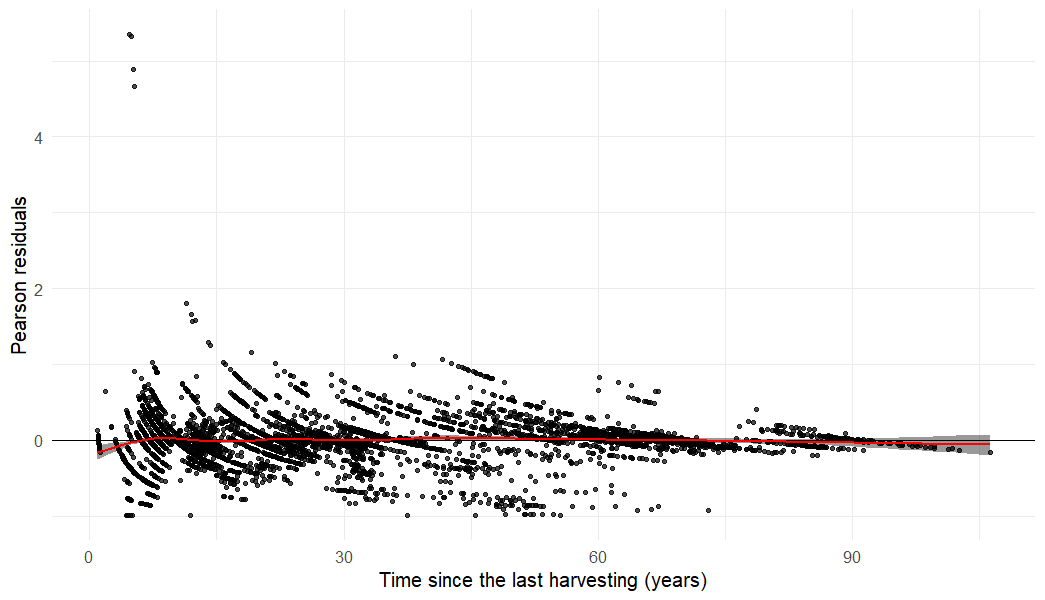

Fig. S6.2: Distribution of the values of the correlation coefficient r squared over the 500 simulations of cross validation. The r squared varied between 0.756 and 0.900.

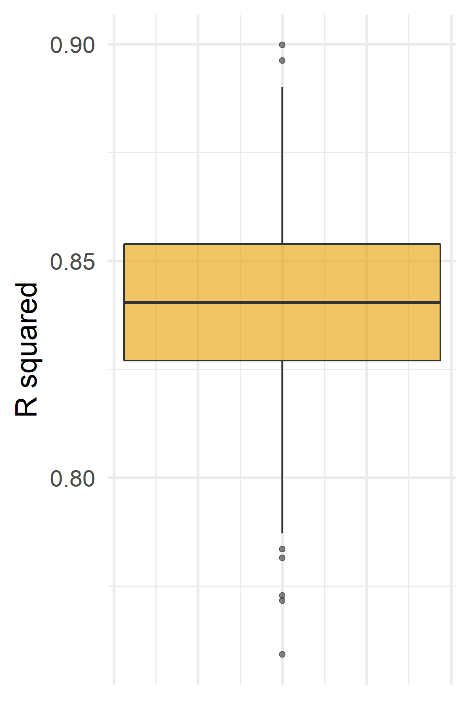

Fig. S6. 3: Estimate variation (mean and standard deviation) after 500 cross validation simulations.
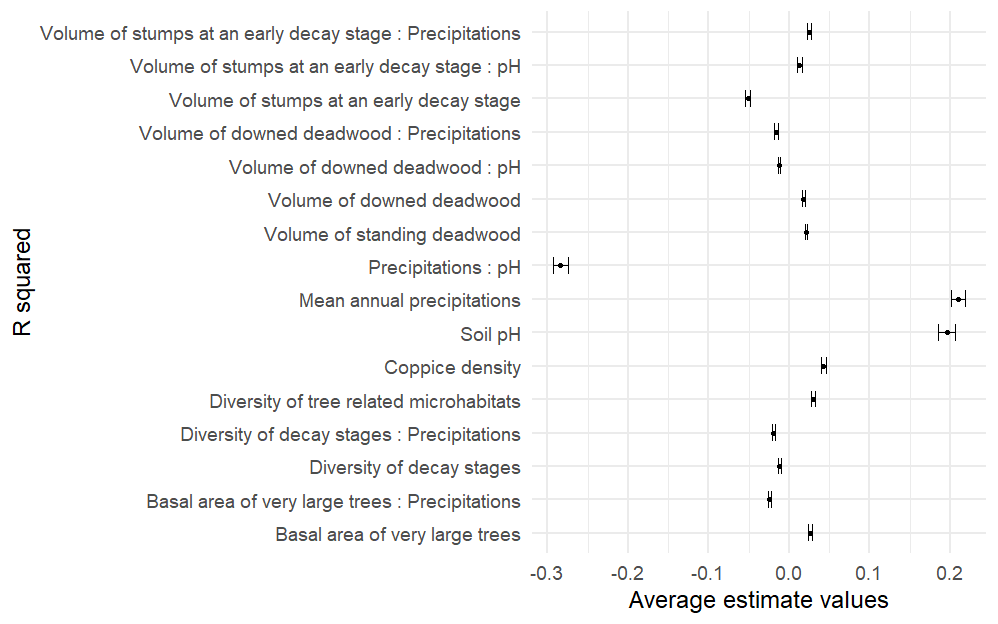

Table S6.1: Standardised estimates, standard errors and p-values rounded to three decimal places for the variables from the simplified generalised linear mixed model with a gamma distribution, log link and “site” random effect explaining the time since the last harvesting of French forest plots. ‘***’ < 0.001, ‘**’ < 0.01, ‘*’ < 0.05, ‘(*)’ <0.1. Volumes and basal areas are all per hectare values.

|  |  | *Standardized estimates (*$\boldsymbol{\beta)}$ | *Standard error* | *p-value (p)* |
| --- | --- | --- | --- | --- |
|  | Intercept | 3.172 | 0.095 | < 2e-16 *** |
| Structural features | Coppice density | 0.052 | 0.006 | 8.71e-8 *** |
|  | Volume of standing deadwood | 0.024 | 0.006 | 1.36e-4 *** |
|  | Volume of large downed deadwood | 0.020 | 0.006 | 1.48e-3 *** |
|  | Basal area of very large trees | 0.029 | 0.008 | 2.89e-04 *** |
| Envrionmental covariates | Mean annual precipitations | 0.244 | 0.040 | 1.17e-09 *** |
|  | Soil pH | 0.191 | 0.049 | 9.08e-05 *** |
| Interactions | Large downed deadwood : Precipitations | -0.010 | 0.006 | 7.98e-2(*) |
|  | Basal area of very large trees : Precipitations | -0.025 | 0.007 | 2.65e-04 *** |
|  | Coppice density : pH | -0.018 | 0.011 | 9.59e-2 (*) |
|  | Precipitations : pH | -0.324 | 0.034 | < 2e-16 *** |
